# Characterization of skin- and intestine microbial communities in migrating high Arctic lake whitefish and cisco

**DOI:** 10.1101/2023.03.08.531621

**Authors:** Erin F. Hamilton, Collin L. Juurakko, Katja Engel, Peter van C. de Groot, John M. Casselman, Charles W. Greer, Josh D. Neufeld, Virginia K. Walker

## Abstract

At high latitudes, lake whitefish (*Coregonus clupeaformis*) and others in the closely related *Coregonus* species complex (CSC), including cisco (*C. autumnalis* and *C. sardinella*), can be diadromous, seasonally transitioning between freshwater lakes and the Arctic Ocean. CSC skin- and intestine microbiomes were collected, facilitated by Inuit fishers at sites on and around King William Island, Nunavut, at the northern range limits of lake whitefish. Community composition was explored using 16S rRNA gene sequencing, with significant differences in microbiota dispersions depending on fishing site salinity for lake whitefish intestine and skin, as well as cisco skin. Overall, lake whitefish intestine communities appeared more variable than cisco and had higher Shannon diversity, suggesting that lake whitefish and their microbiomes could be more susceptible to environmental stress possibly leading to dysbiosis. Although cisco condition was similar among distinct seasonal habitats, the higher average lake whitefish condition in freshwater rivers suggests that fishing these diadromous whitefish in estuaries may be optimal from a sustainable fishery perspective. Taken together, the impact of changing habitats on fish condition and different microbial composition may inform new approaches to CSC health in fisheries and aquaculture, in addition to being relevant for northern Indigenous peoples with subsistence and economic interests in these resources.

## Introduction

Lake whitefish (*Coregonus. clupeaformis*) and cisco (including Arctic cisco, *C. autumnalis,* and sardine cisco, *C. sardinella*), are salmonids that include such similar species that they are often referred to as being part of the *Coregonus* species complex or CSC (Scott and Crossman, 1973). Lake whitefish are commonly found in freshwater lakes in southern latitudes, but they, in addition to cisco, have been reported in northern latitudes within the Yukon River and James-Hudson Bay watersheds (Morin *et al*., 1982; Brown *et al*., 2007). Hybridization has been recorded between each of these three CSC fishes in regions where their ranges overlap such as the ocean basins of Chantrey Inlet and Rasmussen Basin within the Kitikmeot region of Nunavut, Canada (Reist *et al*., 1992; Driver, 2019). Indigenous Knowledge or Inuit Qaujimajatuqangit (IQ) from community fishers in the nearby hamlet of Gjoa Haven teaches that CSC follow the annual sea migration of another salmonid, Arctic char (*Salvelinus alpinus*). Thus, in this region lake whitefish have adopted a diadromous life history similar to cisco and Arctic char. It is therefore of considerable interest to compare these Arctic lake whitefish at the northern limit of their range to their close cisco relatives for physical condition as well as their host-associated microbiota, as they cross salinity and seasonal gradients during migration.

As indicated, lake whitefish, belying their name, may increase fitness by adopting a diadromous life strategy, possibly in response to the relatively low productivity of oligotrophic Arctic lakes (Finstad and Hein, 2012). The physiological challenge required to cope with this life history trait at northern latitudes could be facilitated by the brackish properties of Arctic seawater bodies relative to the Atlantic and Pacific Oceans (Serreze *et al*., 2006; Carmack, 2007). In salmonids, appropriate osmoregulation during the transition between freshwater and saline environments demands behavioural modifications as well as changes to the gills, alimentary tract, epithelium, and kidney function (McCormick, 2012; Kononova *et al*., 2019). Immune systems must remain robust during these transitions and microbial communities associated with fish mucosal surfaces play an important role in immune system maintenance and fish health (Llewellyn *et al*., 2014). As a result, changes in fish microbiota across environmental gradients could result in a risk for dysbiosis, with a corresponding threat of host disease.

In Atlantic salmon (*Salmo salar*), experimental salinity changes between fresh- and saline-waters resulted in a turnover of bacteria associated with the skin and intestine (Dehler *et al*., 2017; Lokesh and Kiron, 2016). Similarly, wild diadromous *S. alpinus* and *C. clupeaformis*, although not as well understood in this respect as Atlantic salmon, exhibit seasonal microbial changes in the intestine microbiota (Element *et al*., 2020a,b). To augment this body of research, we hypothesized that CSC would exhibit shifts in both skin and intestine microbiota during seasonal migration as they follow the Arctic char run in the Kitikmeot region. Accordingly, lake whitefish and cisco were netted from IQ-informed fishing sites along annual runs in autumn, as well as under the ice of lakes where the fish overwinter, and their respective skin and intestinal samples were assessed using high-throughput sequencing of amplified 16S rRNA genes.

Furthermore, core microbiomes, taxa which likely provide essential functions to the fish, were determined within and between seasonal habitats (Lloyd-Price *et al*., 2016), in addition to assessing the general health of the fish as calculated by condition (K factor, a measure of length to weight). Since, as noted, the sampled lake whitefish are from the northern extent of their range (Scott and Crossman, 1973), CSC microbial community composition and physical measures should further aid our understanding of fish microbiota and their response to environmental stress (Rocca *et al*., 2019). In addition, by extension, it is hoped that this analysis may inform stock health, and provide data for the further development of sustainable lake whitefish fishing practices in Arctic communities.

## Methods

### Sample collection

Fishing was conducted in the western Canadian Arctic, including rivers, lakes and estuaries adjacent to the ocean basins of Chantrey Inlet and Rasmussen Basin within the Kitikmeot region of Nunavut. Sampling was done at traditional Inuit fishing sites both on King William Island and south toward the mainland within the inlet of Back River and east across Rasmussen Basin to the Murchison River, as well as saline water sites Back House Point and Legendary River, with closely adjacent sites grouped (Table 1). Fishing locations were selected based on IQ sharing with local Inuit elders and in association with the Hunters and Trappers Association (HTA) of Gjoa Haven, Nunavut. Sites were further designated as autumn-saline, where lake whitefish and cisco were sampled at river estuaries as they migrated from the ocean, autumn-fresh, where fish were sampled along rivers as they returned to lakes, and winter-fresh, where fish were netted during their residency under the ice. Fish were catalogued and weighed, fork length (mm) was measured, and dissected otoliths were aged as previously described (Campana *et al*. 2008; Casselman *et al*., 2019). All samples were given a unique barcode linked to each fish (Wu *et al*., 2020). Fish skin and intestine samples were aseptically collected from fish caught in commercial and subsistence fishing nets or occasionally on fish spears, frozen at - 20 °C then shipped frozen to the laboratory, as previously described (Hamilton *et al*., 2019).

**Table 1.** Fishing site numbers and map names of the sample locations with GPS coordinates in decimal degrees and the salinity of the water collected from those sites as indicated.

| Site | Site Name <sup>1</sup> | Season(s) | Water Type | Conductivity (µScm<sup>-1</sup>) <sup>2</sup> | Latitude | Longitude |
| --- | --- | --- | --- | --- | --- | --- |
| 1 | Port Parry, KWI | Winter | Fresh | 286 | 69.56 | -97.44 |
| 6 | Back House Point | Autumn | Saline | 8 240 | 67.46 | -95.36 |
| 7 | Legendary River | Autumn | Saline | 3 450 | 67.52 | -96.44 |
| 8 | Murchison River | Autumn | Fresh | 225 | 68.57 | -93.38 |
| 10 | Merilik Lake, KWI | Winter | Fresh | NA | 68.57 | -97.33 |
| 13 | Back River | Autumn | Fresh | 19 | 66.96 | -95.30 |
| 18 | Back River (above Hayes River) | Autumn | Fresh | NA | 67.15 | -95.36 |
| 19 | Back River (below Hayes River) | Autumn | Fresh | NA | 67.11 | -95.31 |

<sup>1</sup> Fishing sites on King William Island (KWI) are indicated with all others situated on the mainland.
<sup>2</sup> Specific conductance was provided for sites where applicable (see Methods), while other sites were designated based on location sampled.

Water samples were taken, if possible, from fishing locations (sites 1, 6, 7, 8 and 13; Table 1). Approximately 2 L of water was filtered through sterile 0.22 μm filters (Pall Corporation, Mississauga, ON, Canada) in triplicate, shipped frozen at -20 °C, and subsequently stored at −80 °C until further processing. Unfiltered water samples (50 mL) were collected in plastic tubes, frozen at -20 °C before being transported to the lab, and then stored at -80 °C. Unfiltered water samples were thawed and assessed for specific conductivity using a conductivity meter (Thermo Fisher Scientific, Waltham, MA, USA). Licences to fish for scientific purposes were obtained in accordance with section 52 of the general fishery regulations of the Fisheries Act, Department of Fisheries and Oceans Canada (DFO). Animal care permits were issued by the Freshwater Institute Animal Care Committee of DFO (S-18/19-1045-NU and FWI-ACC AUP-2018-63).

### DNA extraction, sequencing, and bioinformatics

DNA extraction and sequencing was performed as described previously (Hamilton *et al*., 2019; Element *et al*., 2020a,b). Skin and intestinal samples were handled aseptically and skin-associated DNA was extracted using a NucleoSpin Soil Extraction Kit (Machery-Nagel GmbH, Düren, Germany) while the intestine-associated DNA was extracted using MoBio DNA Extraction Kit (QIAGEN, Hilden, Germany) both as described (Hamilton *et al*., 2019; Element *et al*., 2020). It is important to note that for these experiments, we did not separate the parasites from the gut samples prior to extraction of DNA, as has been recommended by some researchers, and thus the DNA from the intestine also includes the communities originating from the parasite microbiome (Kashinskaya *et al*., 2020). DNA from water samples was extracted using hexadecyl trimethyl ammonium bromide and phenyl/chloroform/isoamyl alcohol as previously detailed (Tremblay *et al*., 2017). For skin and intestinal samples, a pre-amplification step targeting the bacterial 16S rRNA V1-V9 region was followed by the amplification of the V4-V5 region using primers 515F-Y (Parada *et al*., 2016) and 926R (Quince *et al*. 2011) as described previously (Hamilton *et al*., 2019; Element *et al*., 2020a). The tissue-derived Illumina libraries were sequenced using a 2 x 250 cycle MiSeq Reagent Kit v2 on a MiSeq platform (Illumina Inc., San Diego, CA, USA). The 16S rRNA gene V4-V5 region was amplified from each water sample and sequenced using the Illumina MiSeq platform (Illumina Inc., San Diego, CA, USA) (Tremblay *et al*., 2017; Cobanli *et al*., 2022). A total of 564 samples comprising water, intestine, and skin as well as controls were analyzed using Quantitative Insights Into Microbial Ecology 2 (QIIME2) (version 2020.6) (Bolyen *et al*., 2019) managed by automated exploration of microbial diversity (AXIOME3) (Min *et al*., 2021). DADA2 (version 2020.6; Callahan *et al*., 2016) was used to remove primer sequences and chimeras, dereplicate and denoise reads. Taxonomy was assigned to amplicon sequence variants (ASVs) using a naive Bayesian classifier pre-trained with the SILVA database (version 138) (Pruesse *et al*., 2007). The prevalence method in Decontam (Davis *et al*., 2018) was used to identify contaminants using a threshold of 0.1 (Table S1). AXIOME3 was used to generate triplots and PCoA ordination using a Bray-Curtis dissimilarity matrix. Skin, intestine, and water sequences are available in the European Bioinformatics Institute (EBI) database under accession number PRJEB48811.

### Age determination, growth estimation, and condition factor

To assess the growth of sampled CSC, mean annual incremental growth curves were calculated for each of the fish species groups. Of the 344 fish (both skin and intestine samples), age data from otoliths was available for 186 lake whitefish and 150 cisco. By extrapolation using a key generated by the size-at-age relationship (Isermann and Knight, 2005), the addition of four lake whitefish and two cisco, resulted in age data for 152 cisco and 192 lake whitefish. Incremental growth curves were calculated by dividing the fork-length (FL; in mm) by age, which provided mean fork length at age (indicated as FL·Age^−1^) for lake whitefish aged 4-43 years (outliers at ages 47, 49, and 56 were excluded for a total of 184) and cisco aged 2-27 (excluding an outlier at age 47 for a total of 135). Standards were produced by plotting the log_10_ of both age and FL·Age^-1^, with subsequent regression analysis to produce an incremental growth standard, taking the antilog of the projected values and plotting a standard incremental annual growth curve. Individual FL·Age^-1^ values for the lake whitefish and cisco were then plotted against this standard curve. In addition, deviation from the growth standard was calculated as relative difference in percent using the following equation:

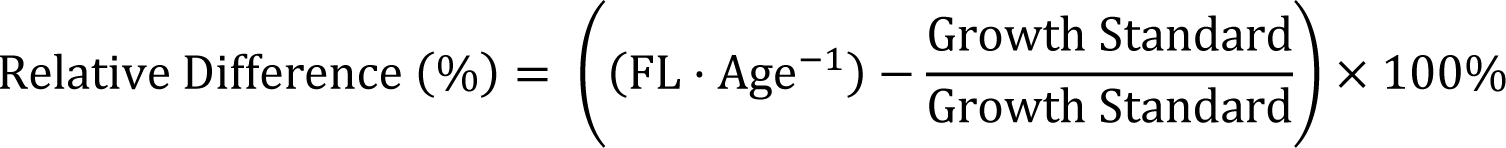

Where the Growth Standard is the value of FL·Age^-1^ predicted by the calculated mean annual incremental growth curve at the specified age of the fish in which the relative difference was calculated. The relative health of individual fish was calculated using Fulton’s condition factor (K) for salmonids (Barnham and Baxter,1998) as:

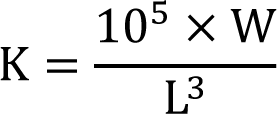

where W is measured in g and L in mm.

### Statistical analysis, data availability and efforts to reduce environmental impact

Significance of group separations was tested by permutational multivariate analysis of variance (PERMANOVA) (Anderson, 2001;2006; Anderson *et al*., 2006) applied using the adonis2 function from the R vegan package (McArdle and Anderson, 2001). In addition, the Shannon index (H) was used to assess species evenness based on the ASV table. Both one-way ANOVAs followed by post-hoc Tukey’s honest significant difference (HSD) tests to correct for multiple testing were performed to assess differences in means between factor groups (using a 95% confidence threshold). Differential abundance analysis between intestine or skin samples from saline- and freshwater habitats was performed using analysis of composition of microbiomes (ANCOM) (Mandal et al. 2015). Pseudocounts were added using low-abundant features (less than 10 total observations) and features unique to less than 5 samples were omitted. Differential abundance testing was performed in QIIME2 and bubble plots were generated using the ggplot2 (Wickham 2009) in R (version 4.0.1).

As well as database submission of DNA sequences (see above), fish samples, parasites, and otoliths have been archived for future access and fish metadata is available in the Polar Data Catalogue (PDC) as open access (PDC#312992; NA profile of IOS 19115:2003, uploaded 5 February 2020, doi.org/10.21963/12992). Measures were taken to reduce the environmental impact of the research activities, including employing community youth to prepare samples, coordination of multiple research projects enabling researchers to volunteer for social science projects, purchase of bulk reagents to share between research groups, making sampled fish available for community food-sharing programs.

## Results

### Characterization of skin and intestine microbiomes of lake whitefish and cisco

The V3-V4 region of the 16S rRNA genes from skin (from 141 lake whitefish and 68 cisco) and intestine (165 lake whitefish and 92 cisco) and the V4-V5 region of the water (30) samples were sequenced. These generated 36,756 unique ASVs across all sample types. A total of 55 skin and 17 intestinal ASVs were identified as contaminants after reference to control or blank amplification reactions using Decontam (Tables S1-S3; Figure S1) and were removed from the dataset before subsequent processing. The 16S rRNA gene profiles showed distinct differences with respect to the most abundant taxa in the two different tissues (Figure S2 and Figure S3).

Skin microbial communities differed when CSC were caught at either ocean or freshwater fishing sites. For the lake whitefish skin microbiota, PCoA ordination showed distinct groupings between the fresh and saline water environments (Figure 1A and Figure S4), as well as a significant difference between dispersions (*p* = 0.001, PERMANOVA). Similarly, in cisco, the skin microbiota grouped based on salinity (Figure 1C and Figure S4), with a significant difference between dispersions (*p* = 0.001, PERMANOVA). Lake whitefish samples were collected in the autumn from saline and fresh-water sites, however, for cisco, an additional winter sampling effort was successful and as a result, comparisons were possible between autumn-saline, autumn-fresh, and winter-fresh seasonal habitats. Again, PCoA ordination for cisco caught in these three seasons indicated distinct groupings in skin communities based on different salinity (not shown), however, there were no significant differences between dispersions for the two different seasonal freshwater habitats for the cisco skin microbiota (*p* = 0.19, PERMDISP2) although PERMANOVA showed a significant difference between dispersions (*p* < 0.001). To assess alpha diversity in each fish species, Shannon diversity (H) was calculated for each of the seasonal habitats (Figure S5). For lake whitefish, there was no significant difference in Shannon diversity for skin microbiota between the two autumn habitats. In contrast, Shannon diversity was significantly greater (*p* < X, one-way ANOVA) in skin samples from cisco from the autumn-fresh seasonal habitat compared to samples from autumn-saline waters.

**Figure 1.**
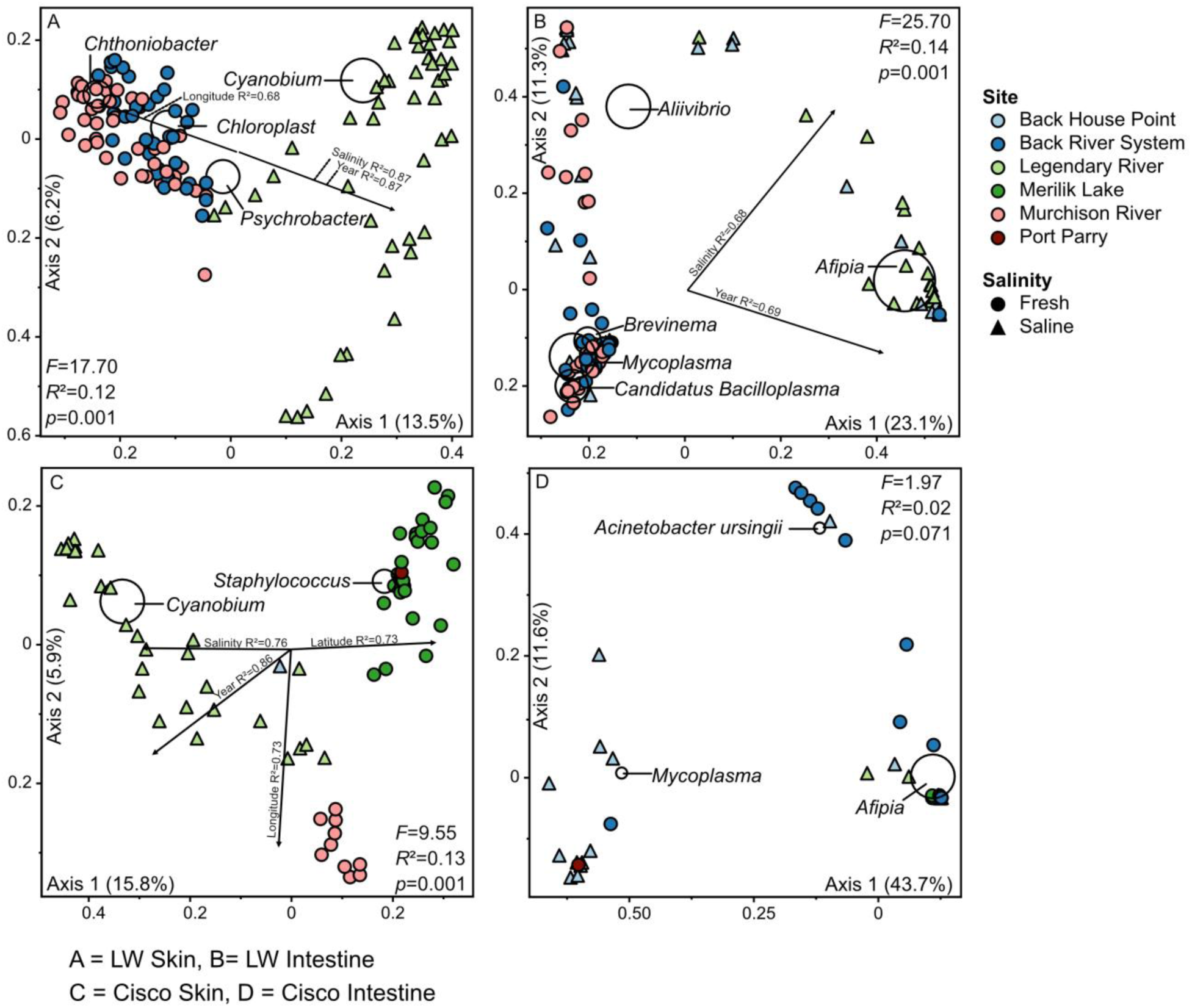
PCoA biplot based on Bray-Curtis dissimilarities showing skin **(A, C)** and intestine **(B, D)** microbiota of lake whitefish (LW; **A, B)** and cisco **(C, D)** at various fishing sites (shown by different colours). PERMANOVA revealed strong significant difference between saline and freshwater dispersions for ordinations A to C. Black circles show ASVs at or above 0.5% **(A to C)** or 0.1% **(D)** relatively abundance across all samples. The lowest taxonomic rank identified with confidence of 0.8 by the classifier is displayed.

Sample-specific differences were observed between seasonal habitat and fishing sites. Generally, lake whitefish skin microbiota in the autumn-saline seasonal habitat was dominated by ASVs affiliated with *Cyanobacteria* (∼21%) and *Psychrobacter* (4%) (Figure 2). Similarly, cisco skin communities sampled at saline sites in the autumn were dominated by *Cyanobacteria* (24-29%) (Figure 3). After swimming upriver, there was a turnover of taxa, but no dominant ASV was identified in the autumn-fresh seasonal habitat in either lake whitefish or cisco skin. Although, as noted, lake whitefish were not caught in the winter, cisco skin microbiota samples from the winter-fresh seasonal habitat communities showed a prevalence of ASVs affiliated with *Candidatus Bacilloplasma* and *Staphylococcus* at both fishing sites (Figure 3).

**Figure 2.**
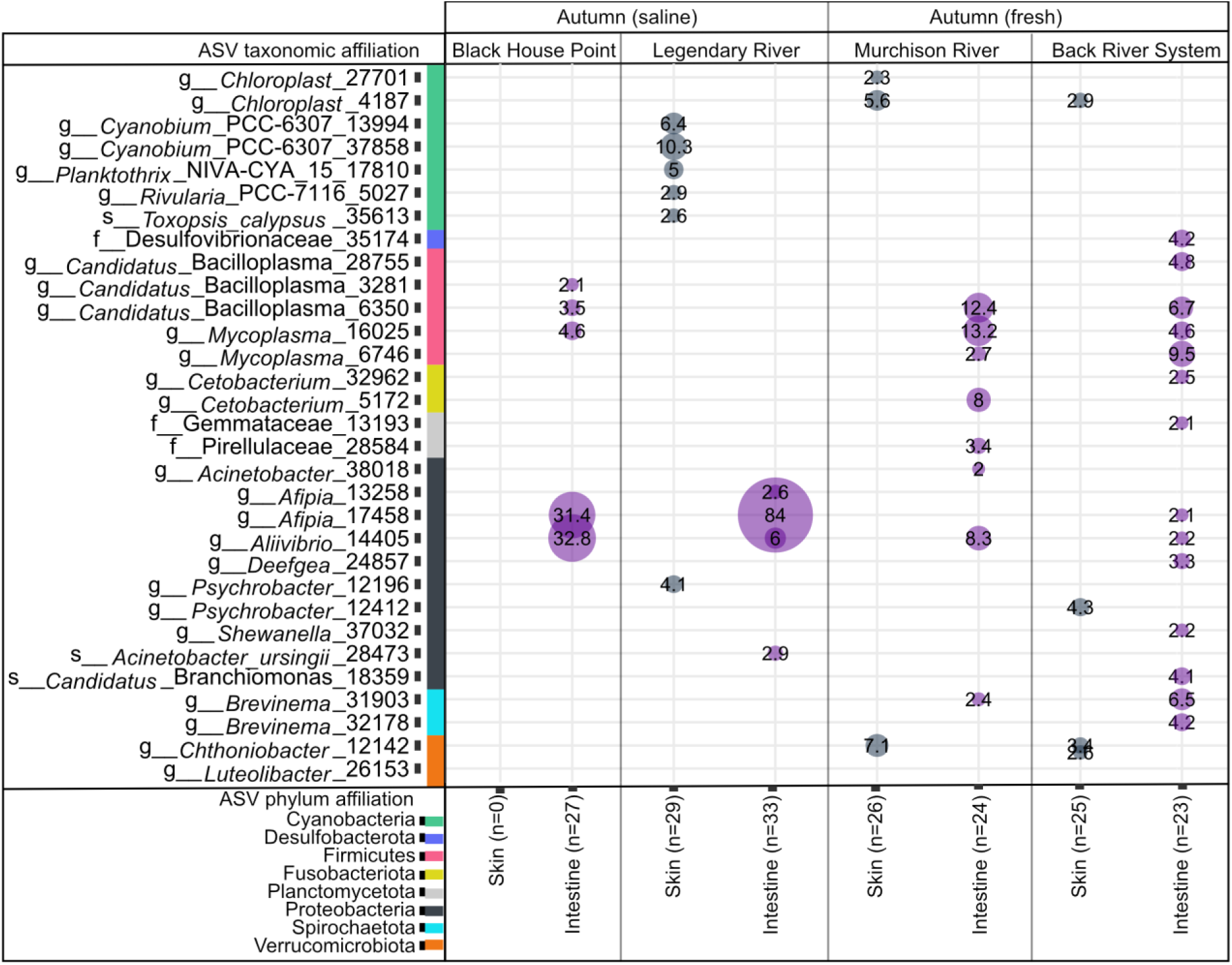
16S rRNA gene profiles for lake whitefish skin and intestine microbiota across autumn-saline and autumn-fresh seasonal habitats and their respective sampling sites. Reads were rarefied to 3,800 and added for each sampling site with total sample numbers shown in the brackets. Only ASVs at or above 2% relative abundance are shown. The lowest taxonomic rank identified with confidence of 0.8 by the classifier is displayed on the Y-axis.

**Figure 3.**
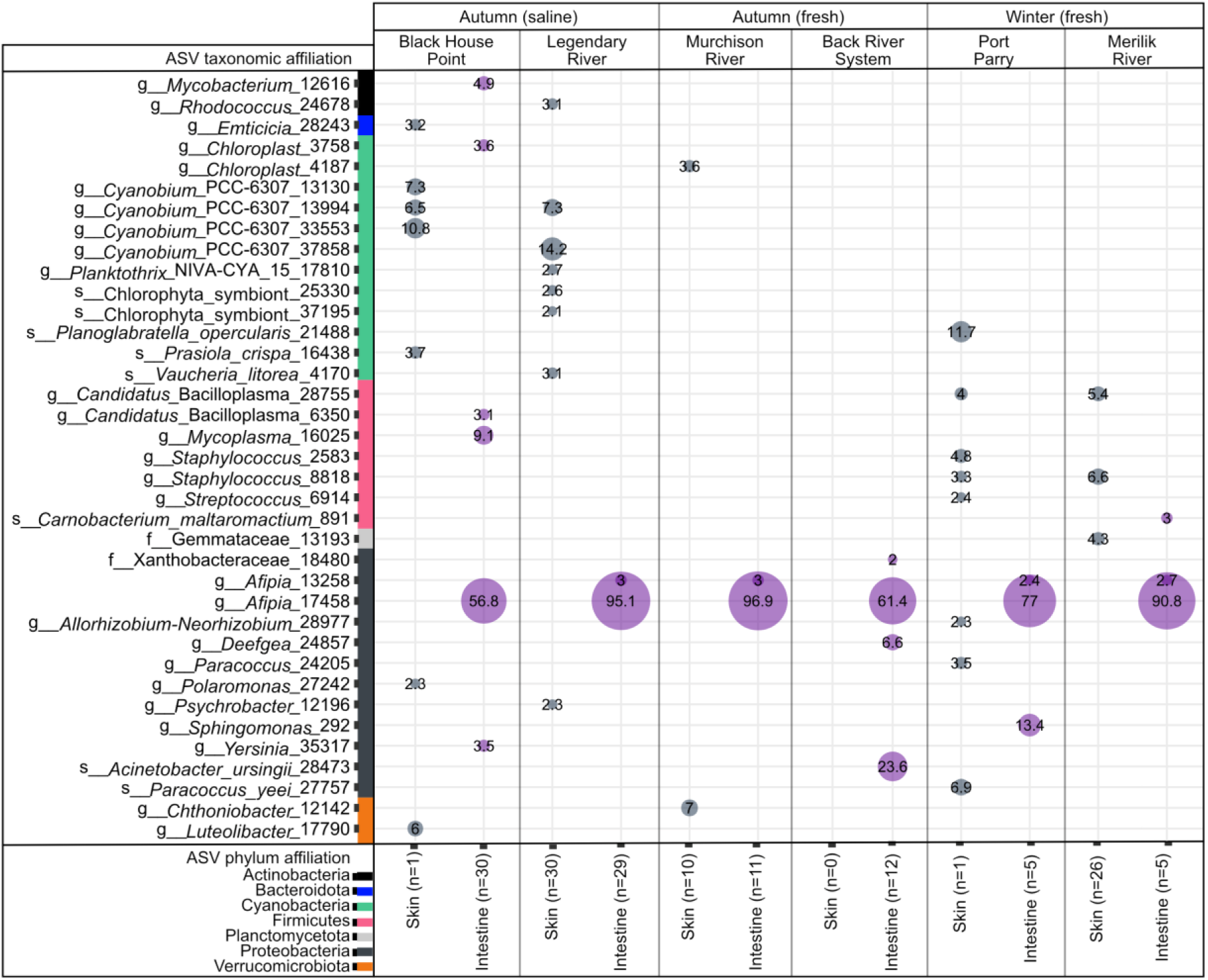
16S rRNA gene profiles for cisco (*C. autumnalis* and *C. sardinella*) skin and intestine communities across autumn-saline, autumn-fresh, and winter-fresh seasonal habitats and their respective sampling sites. Reads were rarefied to 3,800 and added for each sampling site with total sample numbers shown in the brackets. Only ASVs at or above 2% relative abundance are shown. The lowest taxonomic rank identified with confidence of 0.8 by the classifier is displayed on the Y-axis.

Taxa that were similar among seasonal habitats in skin or intestine of lake whitefish and cisco, the “core microbiota”, defined as being present across 50% of samples in each habitat, were identified and the relative percentage contribution of each was calculated. This was best visualized using differential abundance or ANCOM analysis. For autumn-caught lake whitefish, the ocean habitat skin core microbiome was dominated by *Cyanobium* (∼18% of ASVs) and *Planktothrix* (5%), with this community changing to a distinct freshwater community with the most frequent ASVs being *Cthoniobacter* (5%), *Leuteolibacter* (2%), as well as chloroplasts (4%) (Table S4). At the phylum level, differential analysis showed that 63% (29.7/47.1 representing ≥1% of the total) of the ASVs obtained from saline waters were represented by Cyanobacteria, but once in freshwater ASVs from this phylum represented less than 2% of all the ASVs returned (excluding chloroplast sequences) (Figure 4). The most differentially abundant ASVs from freshwater-sampled lake whitefish were assigned to Verrucomicrobiota, but again, the majority of ASVs represented < 1% of the total. Similarly in skin samples from ocean-caught cisco, 75% (29.7/39.6 for those ≥ 1% of the total) of the ASVs were from Cyanobacteria but this taxon was hardly notable in the diverse ASVs from freshwater-caught fish. The most differentially abundant phylum was Firmicutes in this case. Core skin microbiota from cisco in the autumn-saline seasonal habitat was dominated by *Cyanobium* (11% of ASVs), followed by a community turnover in the autumn-fresh habitat where *Candidatus Bacilloplasma* (4%) was found in a majority of samples.

**Figure 4.**
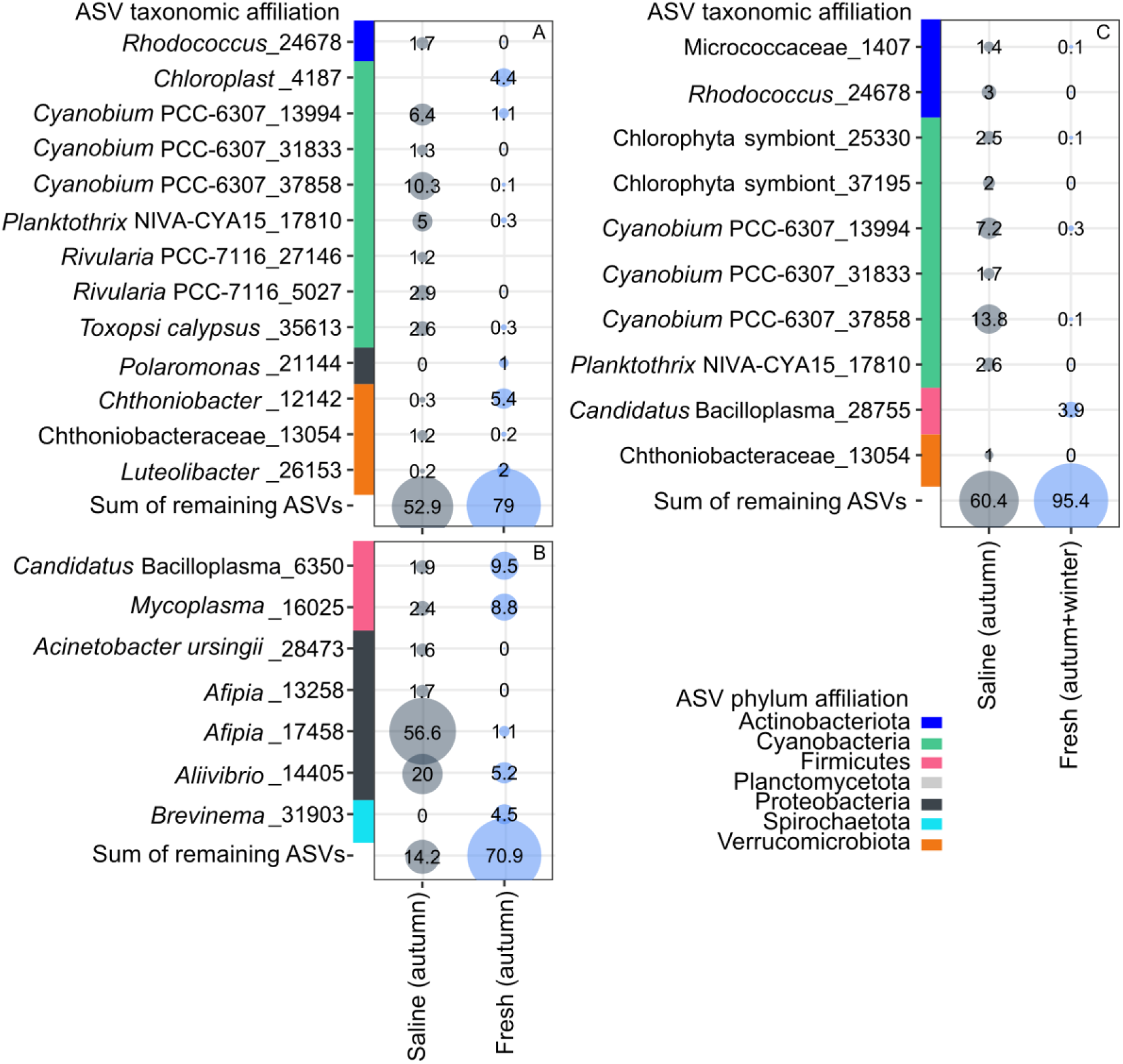
Differential abundance analysis of microbiota from lake whitefish (**A**, **B**) and cisco (**C**) derived from skin (**A, C**) and intestine (**B**). Only ASVs at or above 1% relative abundance within each group of significant ASVs are shown, with the phylum affiliations shown by bubbles of different colours.

PCoA ordination of lake whitefish intestine microbiota showed groupings based on salinity (Figure 1B and Figure S4), with significant differences between dispersions for fresh and saline environments (*p* = 0.001, PERMANOVA). In contrast, cisco intestine microbiota showed no distinct grouping based on salinity and thus no significant differences between freshwater and saline environments (Figure 1D) (*p* = 0.07, PERMANOVA). Intestines from freshwater-caught lake whitefish showed significantly higher (*p* < 0.001, one-way ANOVA) Shannon diversity than those sampled from saline waters (Figure S5). Alpha diversity in cisco intestinal microbiota showed no significant differences between seasonal habitats.

When obtained from the autumn-saline habitat, lake whitefish intestine communities were dominated by *Afipia* and *Aliivibrio* (at 31-84% and 6-33%, of the ASVs, respectively). However, as autumn migration progressed, *Mycoplasma* and *Candidatus Bacilloplasma* together represented ∼25% of the community ASVs, depending on the river (Figure 2). Cisco intestine was comparatively less diverse, with the community dominated by *Afipia* (57-97% of the ASVs) in all seasonal habitats (Figure 3). Core microbiota, visualized by differential abundance or ANCOM analysis showed that in intestines sampled from autumn ocean-caught lake whitefish, the core gut saline-sampled microbiome was dominated by *Afipia* (57% of ASVs) and *Allivibrio* (20%) (Table S4), while the core microbiome in freshwater again was distinct with *Candidatus Bacilloplasma* (10%), *Mycoplasma* (9%) *Brevinema* (4%) and with lesser relative abundance of *Allivibrio* (5%). At the phylum level, there was a turnover from a microbiome dominated (93%; 79.9/85.8 ASVs representing >1% of the total) by Proteobacteria to diverse bacteria with more differentially abundant ASVs from Firmicutes. In contrast, the intestinal core microbiome of cisco persisted across each of the autumn-saline, autumn-fresh, and winter-fresh seasonal habitats, in which *Afipia* was the sole representative across 50% of samples within all of the habitats (Figure 3), and no differential abundant ASVs were detected for cisco intestine samples (not shown).

When Shannon analysis was used to examine distinctions between tissues, lake whitefish skin microbiota showed significantly higher diversity than the intestinal samples (*p* < 0.001, one-way ANOVA) (Figure S5). Similar results were seen for comparisons between skin and intestine microbiota in cisco, where skin microbiota diversity was significantly higher than that from intestines (*p* < 0.001, one-way ANOVA) (Figure S5). Comparison of microbiota between the fish species was also of interest; there was no significant difference in Shannon diversity between lake whitefish and cisco skin microbiota. However, comparisons of intestine microbiota between fish species showed that Shannon diversity was significantly higher (*p* < 0.001, one-way ANOVA) (Figure S5) in lake whitefish.

### Comparison of CSC skin and intestine microbiota to water microbiota

When ordinated based on Bray-Curtis distance, the dispersion of lake whitefish skin microbiota showed a significant difference from that of the water microbiota at corresponding fishing sites (Figure 5A) (*p* = 0.001, PERMANOVA). The dispersion of lake whitefish intestine microbiota was also significantly different from that of the corresponding water microbiota (Figure 5B) (*p* = 0.001, PERMANOVA). Dispersion between cisco skin microbiota and that of the corresponding water also appeared distinct and showed significant differences (Figure 5C) (*p* < 0.001, PERMANOVA), and with obvious dispersion between intestine microbiota and that of corresponding water sites (Figure 5D), with significant differences (*p* = 0.001, PERMANOVA). Thus, fishing site water microbiota and the CSC microbiomes are distinct.

**Figure 5.**
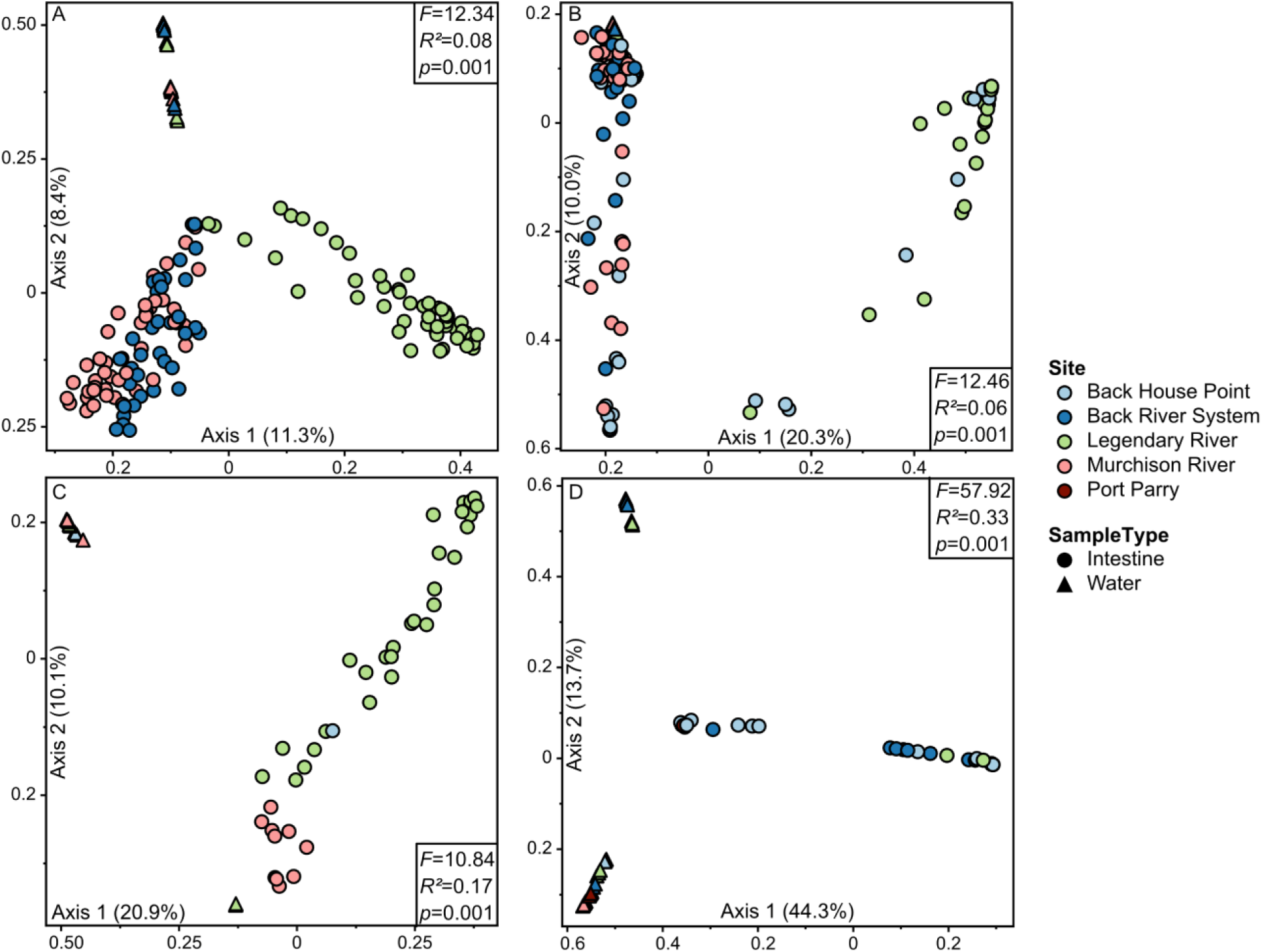
PCoA ordinations based on Bray-Curtis dissimilarities showing microbiota from lake whitefish (**A**, **B**) and cisco (**C**, **D**) associated with skin **(A, C)** and intestine **(B, D)** communities along with water samples from corresponding fishing sites. PERMANOVA revealed strong significant differences between fish microbiota and water samples from corresponding fishing sites.

The differences between the water communities and the CSC microbiomes were explored by the identification of prominent taxa for the different fishing sites (Figure 6). There were differences in the most relatively abundant taxa in different environmental habitats and even from different fishing sites sampled at the same season. At ocean sites south of King William Island, the most abundant ASVs (>2% relative abundance) representing phyla derived from Bacteroidota or Cyanobacteria. The most abundant ASVs from autumn freshwater fishing sites were from Bacteroidota and Proteobacteria from the mainland east of the island or Bacteroidata and Cyanobacteria in the Back River system south of the island. Winter freshwater sampling from under the ice in the winter at fishing sites on the north of the island yielded most relatively abundant ASVs in Actinobacteria and Cyanobacteria. Since the focus of this investigation was not to identify bacteria at different Arctic sites but to investigate if the fish microbiomes simply reflected the water microbiota, an investigation of the water ASVs was not done, but is available in the database. For example, the top two most abundant taxa in saline water samples collected in the autumn at two fishing sites were either taxa from Flavobacteriaceae and Rhodobacteracae (both 4% of ASVs) or *Cyanobium* sp. (5%) and Sporichthyaceae (2%) (Figure 6). In contrast, the most frequent genera at freshwater fishing sites included *Sediminibacterium* sp., previously recovered in Arctic rivers and *Dinobryon* sp., a filamentous plankton known from oligotrophic lakes (Northington *et al*., 2019; Papale *et al*., 2020).

**Figure 6.**
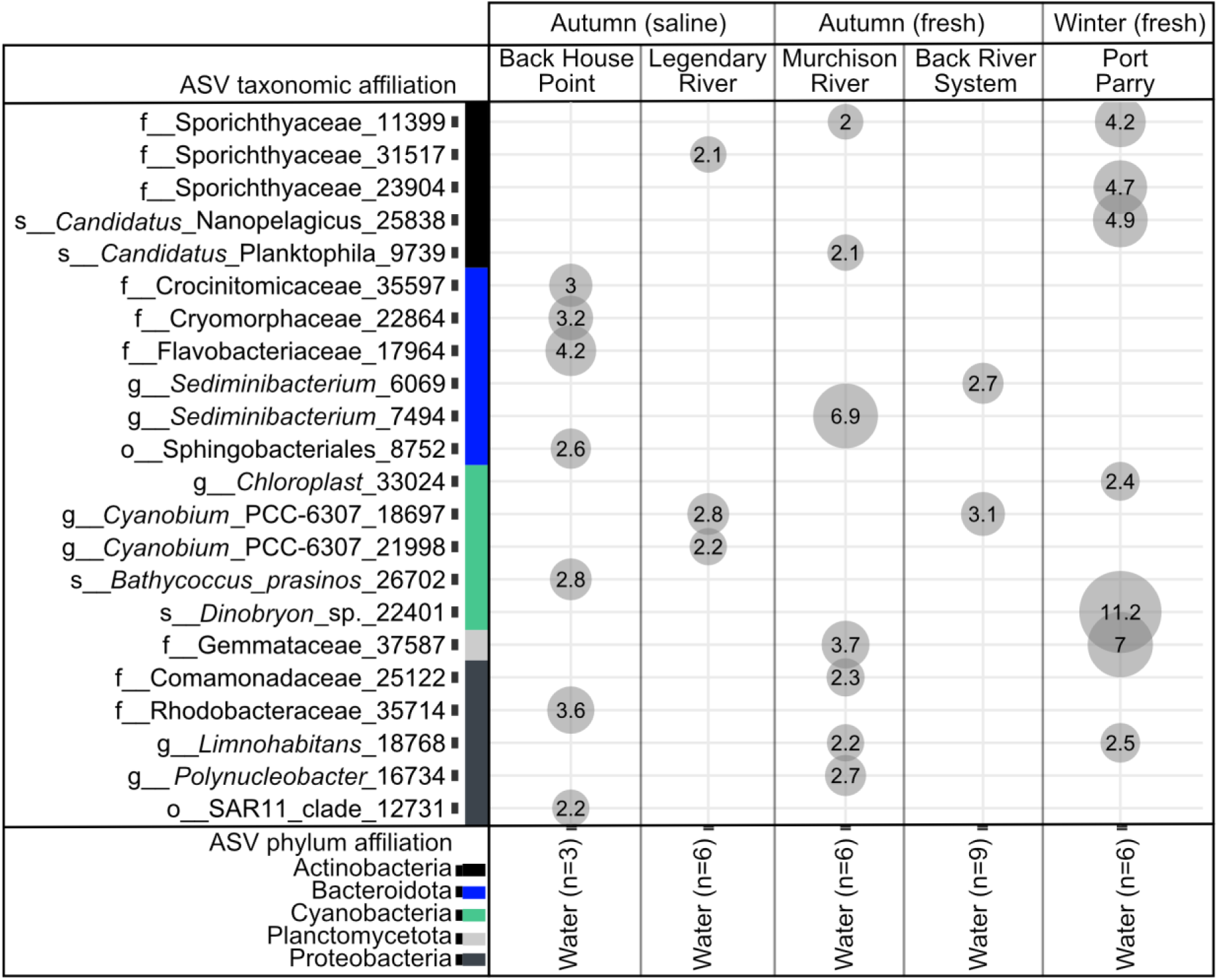
16S rRNA gene profiles for water samples at autumn-saline, autumn-fresh, and winter-fresh seasonal habitats and their respective sampling sites. Reads were rarefied to 3800 and added for each sampling site with total sample numbers shown in the brackets. Only ASVs at or above 2% relative abundance are shown. The lowest taxonomic rank identified with confidence of 0.8 by the classifier is displayed on the Y-axis, with the phylum affiliation indicated by different colours.Tables

### Lake whitefish and cisco mean annual incremental growth at age and condition

To assess the ecology of the sampled lake whitefish and cisco, the mean annual incremental growth (FL · Age^-1^) of the samples was plotted by age against a calculated incremental growth standard curve (Figure S6). The calculated relative percent difference from the standard growth curve resulted in an average deviation of 0.78% (SD=10.5) and 2.46% (SD=15.5), for lake whitefish and cisco, respectively. When lake whitefish growth standard curves were compared, the percent deviation from the growth standard for ocean-netted fish was greater (*p* = 0.032, one-way ANOVA) than those caught in freshwater (Figure S7). In contrast, there were no significant differences seen in cisco caught in different seasonal habitats, nor when further tested based on the factor of salinity (Figure S7B). When condition factor was calculated and compared across seasonal habitats, lake whitefish condition was significantly higher (*p* < 0.001, one-way ANOVA) in the freshwater habitat than in the marine environment, with no significant differences in cisco condition across different seasonal habitats (Figure S8).

## Discussion

### Characterization of CSC microbiota from salt and freshwater fishing sites

Taxa associated with the skin and gut microbiota in migrating lake whitefish caught in waters found on and surrounding King William Island in Nunavut changed across salinity gradients, consistent with the influence of changing environments in other salmon, and the influence of seasonal habitat in migrating Arctic char (Dehler *et al*., 2017; Lokesh and Kiron, 2016; Hamilton *et al*., 2019; Element *et al*., 2020a; Ghosh et al., 2022). As in lake whitefish, some taxa in the skin microbiota of cisco turned over as the fish swam from the sea and moved up-river to overwinter in lakes (Figures 1-3 and Table 1). In contrast, cisco intestinal microbial communities remained relatively constant throughout the autumn and winter seasons.

Independent of seasonal habitat, gut microbiota was dominated by *Afipia,* comprising 57-97% of the taxa, known to oxidize ammonia or nitrite and be associated with biofilms (Roveto and Schuler, 2019). An important influence on the fish gut microbiome is diet (Boutin *et al*., 2013; Uren Webster et al. 2018; 2020). Thus, the constancy of taxa in different environments could suggest that cisco ate little during their migration and subsequent overwintering in the northern oligotrophic lakes, but this would also be true of Arctic char, and their gut microbiota was reported to turnover in the autumn (Element *et al*., 2020a). Alternatively, the dominance of *Afipia* may underscore its importance in gut function and biofilm formation, explaining this taxon’s major contribution to the cisco gut core microbiome irrespective of water salinity.

Although *Afipia* also showed the highest relative abundance in saline-sampled lake whitefish intestines (57% of ASVs), the additional substantive contribution by *Aliivibrio* (20%), an opportunistic taxon linked to unhealthy or diseased salmon in aquaculture facilities, could be of concern (Wang *et al*., 2018; Bozzi *et al*., 2021). During the autumn migration, however, both these two taxa were largely displaced by a more variable community. Considering that cisco and lake whitefish are so closely related that they can hybridize (Reist *et al*., 1992), and that drinking is not required to maintain osmotic balance when entering freshwater, it is striking that the gut communities in lake whitefish, but not cisco, shifted dramatically during migration. Once in freshwater, diverse intestine taxa were identified in lake whitefish, with the most prevalent taxa, *Candidatus Bacilloplasma* and *Mycoplasma,* together representing 19% of the ASVs.

*Mycoplasma* sp. is frequently reported in salmonids and is known to adhere tightly to host epithelium where it likely contributes to biofilm formation (Razin and Jacobs, 1992; Bozzi *et al*., 2021). *Candidatus Bacilloplasma* has been reported in aquatic organisms including fish, crabs and shrimp, and is of possible concern since it is linked to diseases in shrimp (Chen *et al*., 2017; Hou *et al*., 2018; Sun and Xu., 2021).

As noted, most lake whitefish populations, at least in more southern regions, do not routinely migrate to the sea throughout much of their range. However, diadromous behaviour in *C. autumnalis* and *C. sardinella* is common and therefore cisco could be better adapted physiologically to this life history than lake whitefish (Scott and Crossman, 1973). There is an optimal diversity range for resistance to dysbiosis (Rocca *et al*., 2019), and we assume that such extents are unique to species and habitat. For example, chum salmon (*Oncorhynchus keta*) show increased microbiome diversity at temperatures that are higher than optimal (Ghosh et al., 2022). It is therefore tempting to speculate that the higher microbial diversity of lake whitefish intestinal microbiota, compared to cisco, reflects the residency of these lake whitefish at the extreme northern extent of their reach. The more variable taxa in lake whitefish then could possibly reflect a relatively precarious state that could make these fish more vulnerable to colonization by potential pathogens, including those identified such as *Candidatus Bacilloplasma* (10% of taxa), associated with diseases of aquatic organisms, *Allivibrio* (5%), linked to diseased salmonids as well as *Brevinema* (4%) and a possible opportunistic pathogen in trout (Brown *et al*., 2019; Bozzi *et al*., 2021; Sun and Xu, 2021). Therefore, we posit that cisco gut populations that are completely dominated by *Afipia* no matter the seasonal habitat, may be protective for dysbiosis. It is noteworthy too that although there is turnover in gut microbiota in diadromous Arctic char, two taxa, *Photobacteria* and *Mycoplasma* anchored the gut microbiota in both seasonal habitats (Element *et al*., 2020a). In contrast, whitefish with their dramatically changing community and relatively abundant taxa of concern, possibly linked to a stressful forced diadromous life history at the northern limit of their range, may be more susceptible to disease.

Like intestinal microbiota, environmental conditions can influence skin communities, including the surrounding waters (Uren Webster 2018; 2020). For the skin microbiota, both CSCs showed near identical core taxa when fished from ocean sites. In both lake whitefish and cisco, Cyanobacteria represented ∼ 25% of the ASVs with the nutrient-cycling, photoautotroph *Cyanobium* sp. (11-18% of ASVs) assigned to the core microbiota in both fish, where this taxon is likely associated with biofilm on the skin (Faria *et al*., 2021). Cyanobacteria also made a major contribution to Arctic char skin communities fished from these same waters (Hamilton *et al*., 2019). However, the addition of *Planktothrix* (5%), a filamentous cyanobacteria to the skin microbiota of lake whitefish but not cisco, may be notable since at least one of these species has shown toxicity to whitefish (Pancrace *et al*., 2017; Ernst *et al*., 2006). As in other fish, taxa in the CSC skin microbiota change when in water with different salinities (*e.g.* Hamilton *et al*., 2019; Sehnal *et al*., 2021). While traveling from the sea, microbiota turned over as indicated by PCoA ordination that showed distinct groupings, and with similar Shannon diversity. Once in the fresh water neither CSC skin microbiome was dominated by particular taxa, but in lake whitefish minor contributors including *Chthoniobacter* and *Leuteolibacter*, with *Candidatus Bacilloplasma* made up the core bacteria. Some of these taxa are known in salmonid or isopod microbiomes or as probable biofilm-formers (Sangwan *et al*., 2004; Kostanjsek *et al*., 2007; Kershaw, 2015; Niu *et al*., 2020).

Microbiota from water samples obtained from CSC fishing sites were not unexpected since the identified taxa have been described previously. What was notable, however, was that there appeared to be less concordance in the most relatively abundant taxa at different fishing sites with similar salinities or collection season, than in each fish species obtained from different sites but with the same seasonal habitat (Figures 2 and 3 *vs.* Figure 6). Water microbiota is thought to strongly influence fish microbiota, presumably as it would bathe the skin communities and be necessarily imbibed as migrating fish enter the ocean (Uren Webster, 2018; 2019).

Consistent with this perception, water microbiota was more diverse than either CSC intestine microbiota and showed higher Shannon diversity than lake whitefish skin microbiota. For all other metrics, CSC skin microbiota showed no significant difference in diversity when compared to the surrounding waters. Thus, microbes in the water appear to more easily colonize skin than the gut, suggestive of a higher selectivity by the fish intestinal tract, compared to the skin. Nevertheless, communities in the different tissues were distinct from those of the respective fishing site water, as well as having significantly different dispersions in ordination, indicating that CSC hosts are colonized by water bacteria but subject to tissue selection. It is worth noting that the dispersions for cisco tissue-associated microbiota and the water communities were particularly striking, again suggesting that lake whitefish might be less selective in orchestrating their microbial assemblages compared to cisco. Although highly speculative, we wonder if host genes that regulate such selectivity in these diadromous lake whitefish at their northern limits may not yet be fixed within their genomes.

### Practical considerations for future fisheries management

In the Arctic, migration of Arctic char and CSC are undoubtedly advantageous due to the higher abundance of prey in the marine environment relative to oligotrophic lakes (Finstad and Hein, 2012). Therefore, it is curious that K, the condition factor of lake whitefish caught in the sea, was significantly lower compared to that in rivers, with this difference not seen in cisco, nor this regions diadromous Arctic char (Element *et al*., 2020a). The lower average condition of ocean-caught lake whitefish cannot be due to morphologically distinct *C. clupeaformis* populations in freshwater rivers and lakes (Solovyer *et al*., 2019) since less than 0.8% of all samples deviated from constructed incremental growth curves. Saline-environment fish were caught at the start of their autumn migration and close to the peak of the Arctic char run in accordance with IQ, which teaches that lake whitefish follow Arctic char upriver. Migration success with its numerous physiological demands, would surely favour fish in high condition, and we therefore speculate that lake whitefish with low condition would be less likely to complete their journey and explain the condition disparity in the two seasonal habitats. This hypothesis may be partially supported by examination of lake whitefish ages plotted by year class that indicate that in some years there may have been less juvenile recruitment, underscoring the costs of migration and the more precarious position of lake whitefish at its northern limits.

Despite the onerous physiological and behavioural challenges of adjusting to differing water salinities, the diadromous lifestyle in this high Arctic region has a fitness benefit, and here we suggest that considerations of such fitness costs are not complete without understanding changes to host microbial communities. Microbiome diversity provides some resistance to dysbiosis, however, as indicated, there is an optimal range of alpha diversity for maximal resistance, such that higher diversity can reveal a community in flux (Rocca *et al*., 2019).

Turnover in community composition results in an increase in potential microbial colonizers including potential pathogens. Cisco, which are well within their optimal range in this region of the Arctic (Scott and Crossman, 1973), showed less diversity in intestine communities composed almost entirely of *Afipia,* and independent of seasonal habitat, compared to their lake whitefish relatives that share the same waters and migration routes. It is therefore possible that the increased diversity in lake whitefish leaves them vulnerable to dysbiosis associated with migration from the sea to the rivers, with commensal taxa acting as opportunistic pathogens, in addition to the identified taxa of possible concern including *Candidatus Bacilloplasma*, *Allivibrio,* and *Brevinema*.

Previous research on Atlantic and chum salmon, Arctic char, and non-migratory lake whitefish have provided fine examples for the study of salmonid microbiomes (*e.g.* Dehler *et al*., 2017; Lokesh and Kiron, 2016; Sevellec *et al*., 2018; Hamilton *et al*., 2019; Solovyev *et al*., 2019; Ghosh et al., 2022), but consistencies in trends seen in diadromous CSC shown here should assist with the consolidation of our understanding of community turnover in a changing environment. From a sustainable fishery perspective, although contrary to routine management advice, netting lake whitefish in estuaries in the autumn may be strategic in that a higher proportion of lake whitefish with low condition that would be less likely to contribute to the next generation would be caught. This then would result in proportionally more fish in the best average condition migrating up rivers to spawn. Characterization of microbiomes of wild-caught fish is also of interest for the growing aquaculture industry (Gomez *et al*., 2013), such that this information could be valuable for future consideration as probiotics in northern aquaculture and, in particular, the proposed lake whitefish farms of the Saugeen Ojibway First Nation (*e.g.* Gobin and Lauzon, 2016; Moniz *et al*., 2021). Taken together, investigation of northern diadromous fish microbiota has and will likely continue to generate practical suggestions for fisheries management and sustainability.

## Acknowledgements

We thank the residents of Gjoa Haven, Nunavut, the Gjoa Haven Hunters and Trappers Association, community fishers, and youth for identifying fishing sites, for their invaluable indigenous knowledge, as well as sampling expertise and preparation. Kristy Moniz is thanked for general laboratory assistance, with age and growth interpretation assistance from Bronte McPhedran, Jordan Balson and Kate Brouwer. We are grateful to Drs. W. Leggett, L. Harris, and K. Hedges, as well as S. Schott and his social science team, for supporting our efforts to investigate Arctic fish.

## Author Contributions

EH, GE and PdeG obtained or prepared samples for sequencing, EH, KE, CG, JC and CJ processed data and assembled figures, VW conceived and designed the study, and all authors contributed to manuscript preparation.

## Supplemental Figures

**Figure S1.**
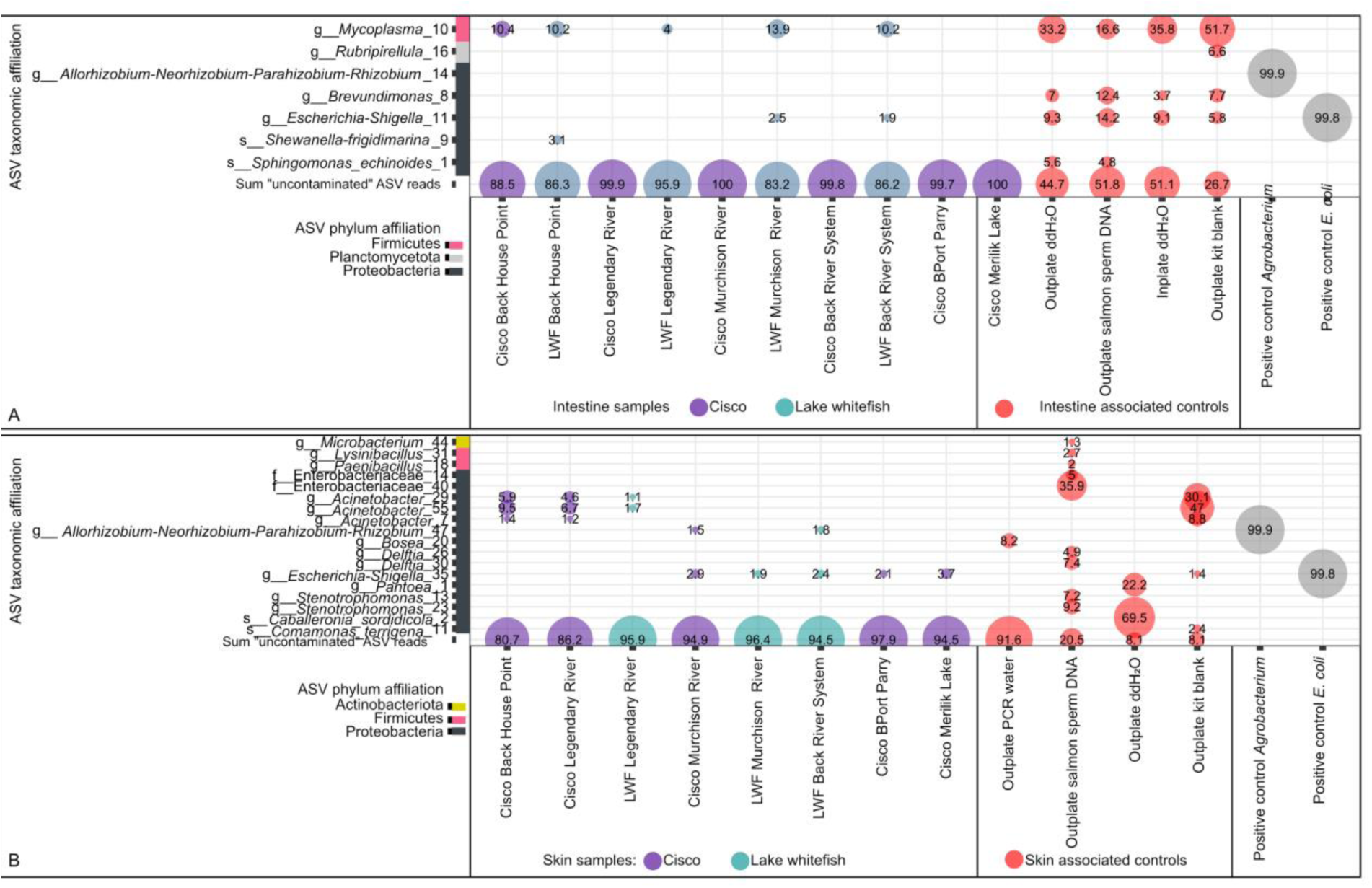
Comparison of contaminant ASVs identified by Decontam in samples and controls. The relative abundance of ASVs is shown for contaminants identified by Decontam for intestine **(A)** and skin **(B)** samples. Only ASVs at or above 1% relative abundance are shown. The sum of all ASVs not identified as contaminants by Decontam are represented as “Sum uncontaminated ASV reads” within each panel. Y-axis label reports lowest taxonomic rank for each ASV with confidence values above the default 0.7 threshold of the classifier. Controls were amplified in single tubes, outside of the 96 well plate (outplate) or within the 96-well plate (inplate). No-template controls (PCR water or autoclaved double distilled water (ddH_2_O), positive controls (*Agrobacterium* or *E. coli* genomic DNA), negative controls (salmon sperm DNA) and DNA extraction kit controls (kit blank, no sample was added) were included.

**Figure S2.**
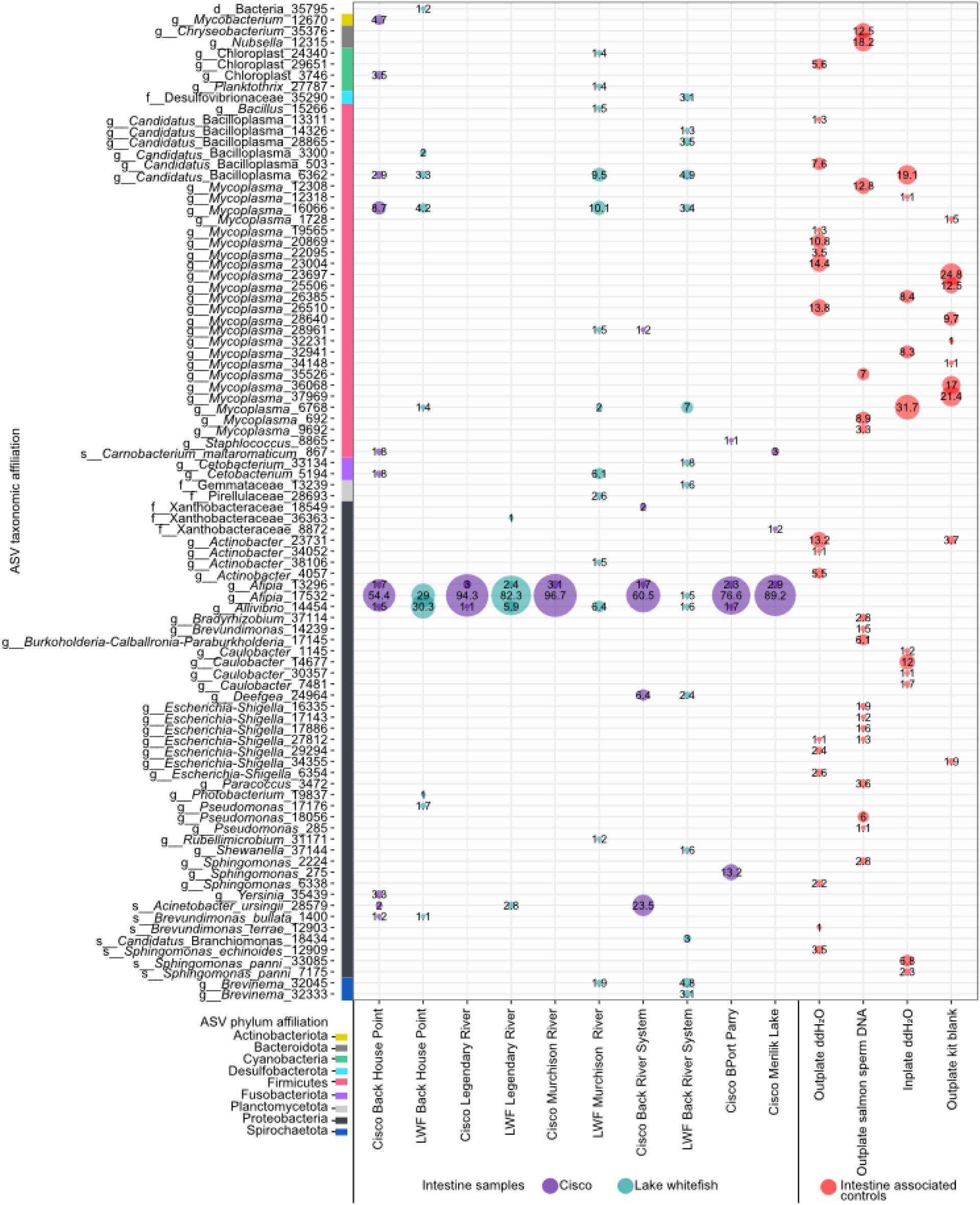
Comparison of 16S rRNA gene profiles in intestine associated samples and controls after removal of ASVs identified by Decontam. Only ASVs at or above 2% relative abundance are shown. Y-axis label reports lowest taxonomic rank for each ASV with confidence values above the default 0.7 threshold of the classifier. Controls were amplified in single tubes, outside of the 96 well plate (outplate) or within the 96-well plate (inplate). No-template controls (PCR water or autoclaved double distilled water (ddH_2_O), negative controls (salmon sperm DNA) and DNA extraction kit controls (kit blank, no sample was added) were included. Replicate samples were merged into one lane.

**Figure S3.**
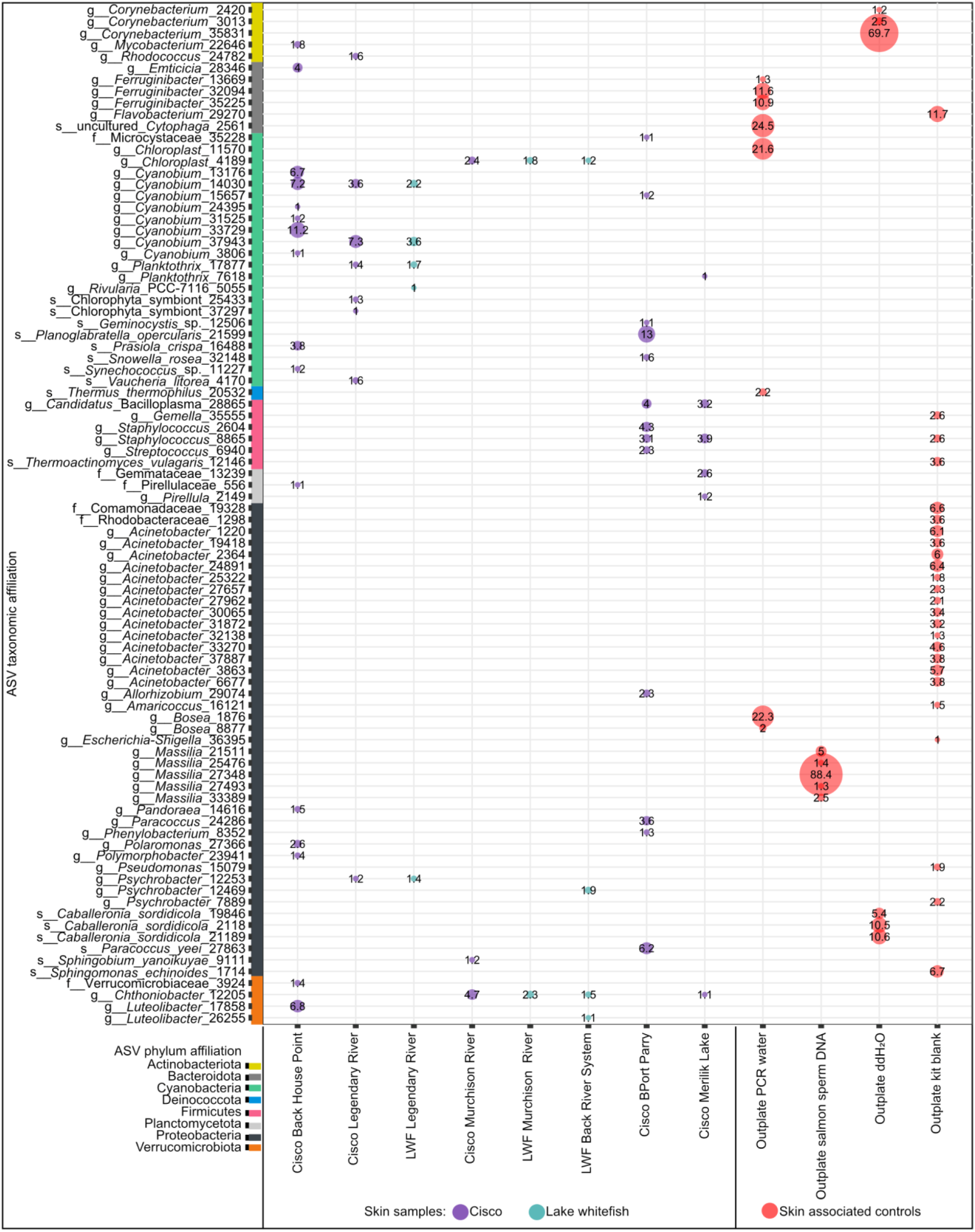
Comparison of 16S rRNA gene profiles in skin samples and controls after removal of ASVs identified by Decontam. Only ASVs at or above 1% relative abundance are shown. Y-axis label reports lowest taxonomic rank for each ASV with confidence values above the default 0.7 threshold of the classifier. Controls were amplified in single tubes, outside of the 96 well plate (outplate) or within the 96-well plate (inplate). No-template controls (PCR water or autoclaved double distilled water (ddH_2_O), negative controls (salmon sperm DNA) and DNA extraction kit controls (kit blank, no sample was added) were included. Replicate samples were merged into one lane.

**Figure S4.**
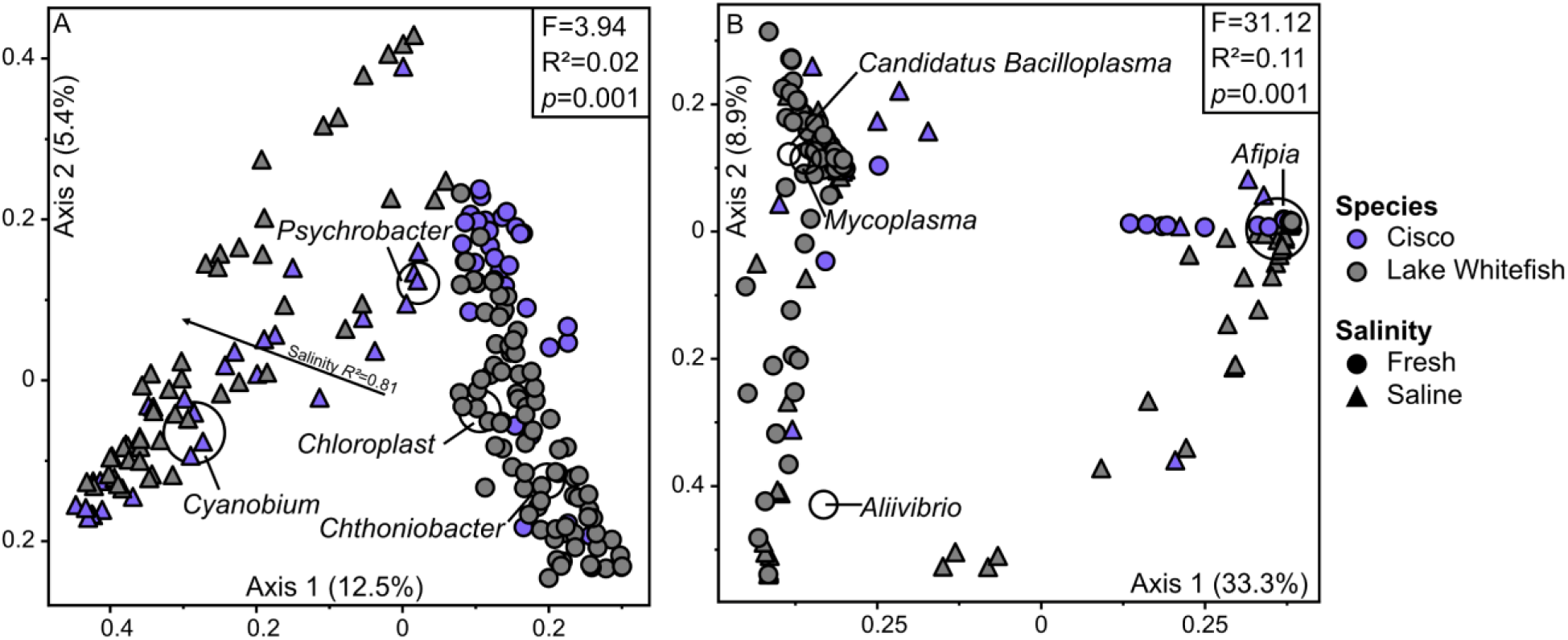
PCoA triplot based on Bray-Curtis dissimilarity showing skin **(A)** and intestine **(B)** microbiota of lake whitefish (*C. clupeaformis*) and cisco (*C. autumnalis* and *C. sardinella*) with 95 fishing sites grouped either as fresh water or saline. PERMANOVA revealed a significant difference between species dispersions. Black circles show ASVs at or above 0.1% relatively abundance across all samples. The lowest taxonomic rank identified with confidence of 0.7 by the classifier is displayed and highlighted by open circles. Samples were rarefied to 4,000 reads.

**Figure S5.**
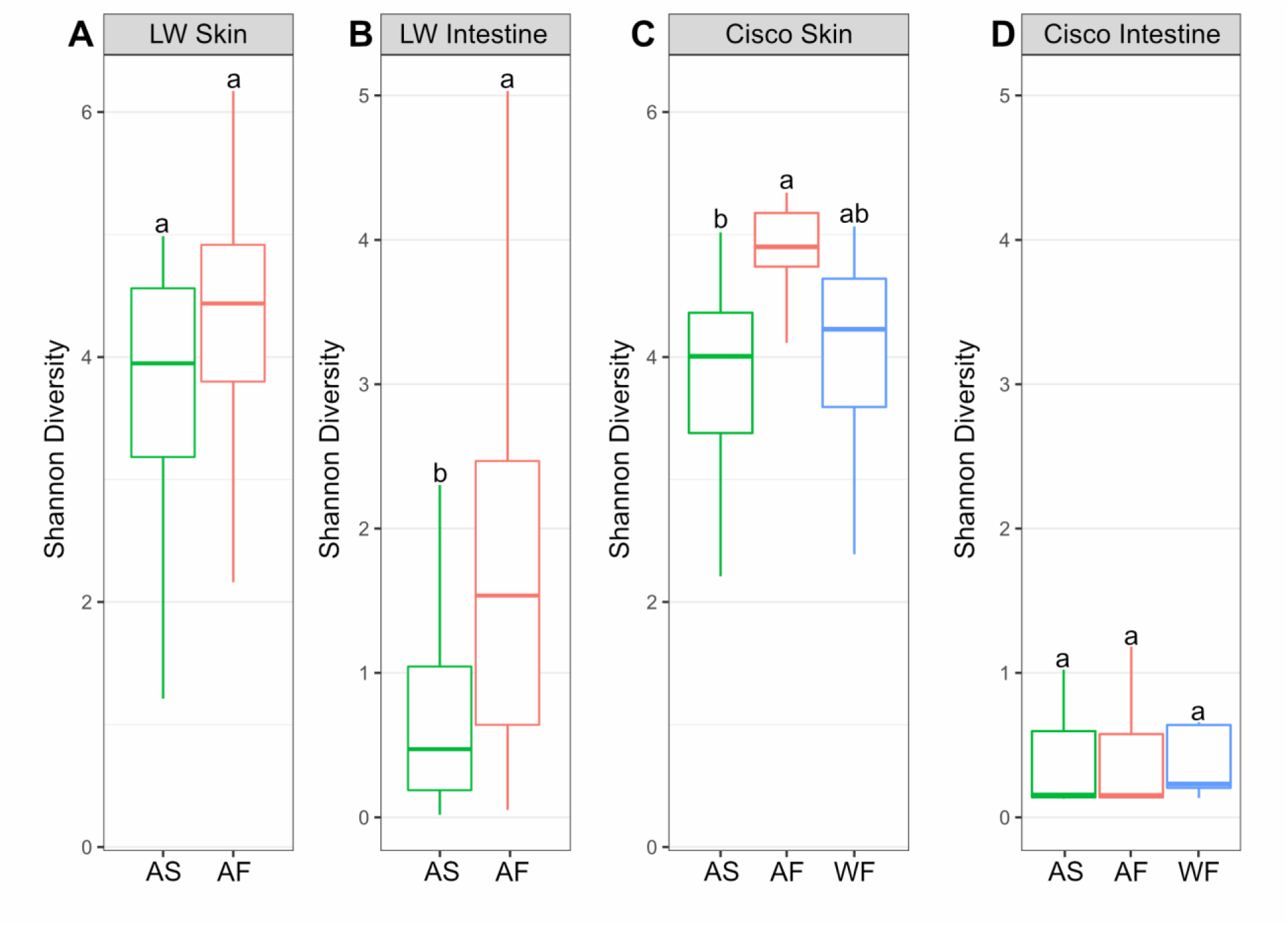
Box plots of alpha diversity measured using the Shannon diversity index metric for variation in seasonal habitats of **(A)** lake whitefish (LW; *Coregonus clupeaformis*) skin across autumn-saline (AS) and autumn-fresh (AF) seasonal habitats, **(B)** lake whitefish intestine across autumn-saline and autumn-fresh seasonal habitats, **(C)** cisco (*Coregonus autumnalis* and *Coregonus sardinella*) skin across autumn-saline, autumn-fresh, and winter-fresh (WF) seasonal habitats, and **(D)** cisco intestine across autumn-saline, autumn-fresh, and winter-fresh seasonal habitats

**Figure S6.**
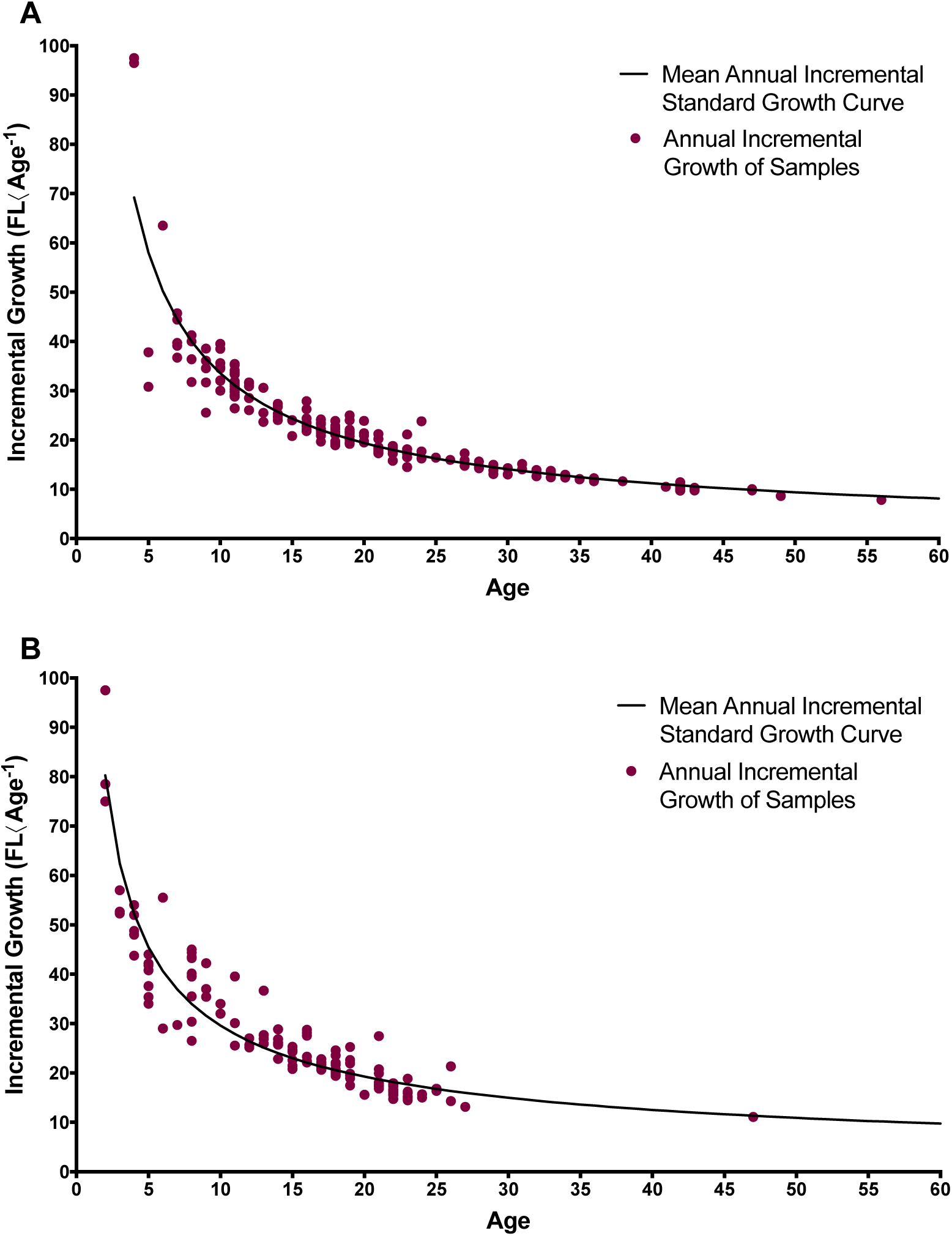
Annual incremental growth at age data against a growth standard curve for lake whitefish and cisco. **A)** Lake whitefish samples (n=188) were plotted against a mean annual incremental standard growth curve. The mean incremental standard growth curve was produced based on whitefish samples aged 4-43 (excluding outliers at ages 47, 49, and 56) and incremental growth at age of each sample was plotted against this curve. **B)** Cisco samples (n=136) were plotted against a mean annual incremental standard growth curve. The mean incremental standard growth curve was produced based on whitefish samples aged 2-27 (excluding outlier at ages 47) and incremental growth at age of each sample was plotted against this curve.

**Figure S7.**
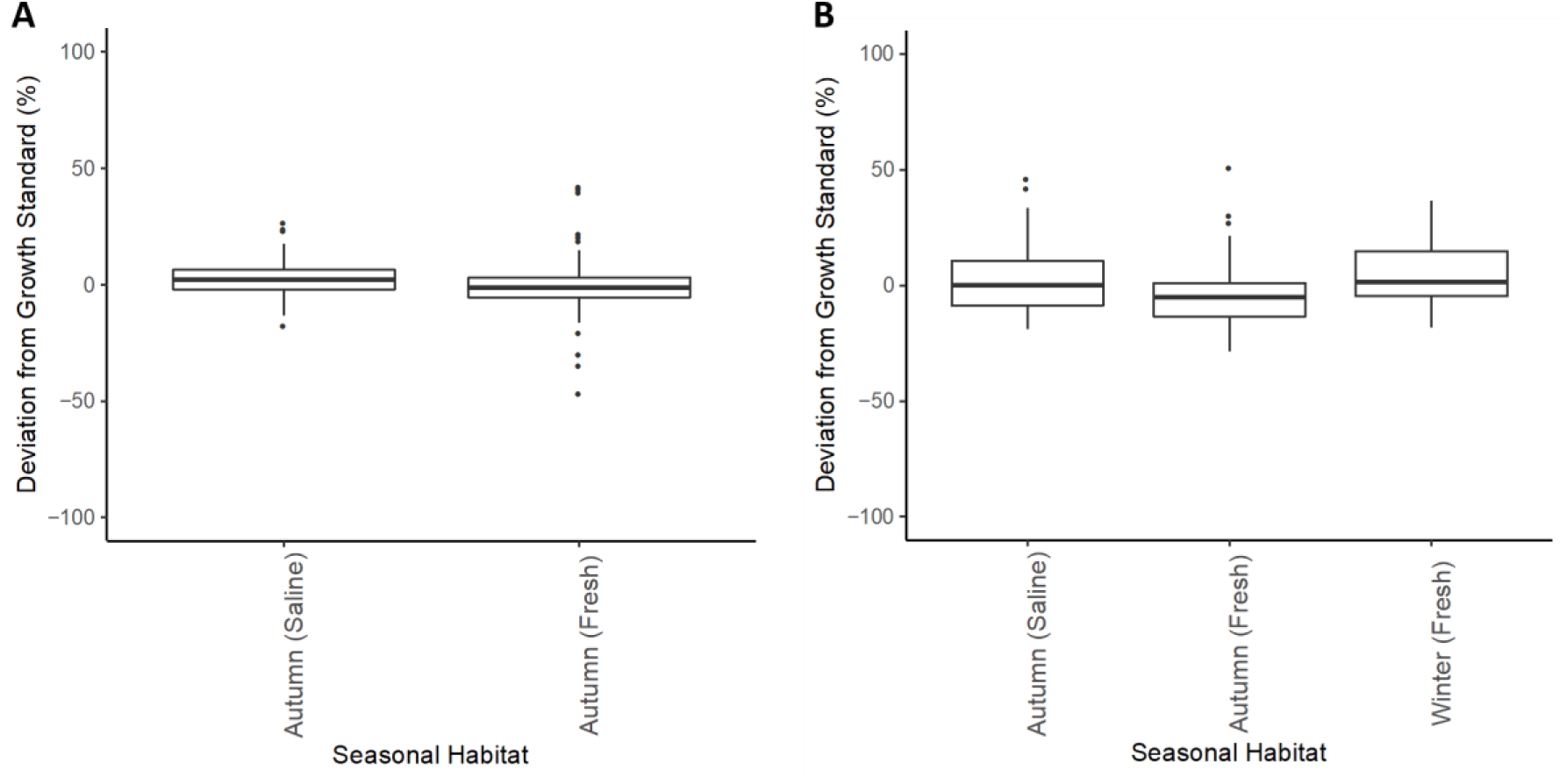
Deviation from standard growth curve, as relative percent difference, with samples categorized based on seasonal habitat, where **A)** represents percent deviation from the standard growth curve for lake whitefish samples (n=188), in which deviation from the growth standard curve is compared between seasonal habitats autumn-saline (n=91) and autumn-fresh (n=97), in which habitats were significantly different (*p* = 0.032, one-way ANOVA) and **B)** represents percent deviation from the standard growth curve for cisco (n=136), in which deviation from the growth standard curve is compared between seasonal habitats autumn-saline (n=63), autumn-fresh (n=31), and winter-fresh (n=42), though there were no significant differences between habitats.

**Figure S8.**
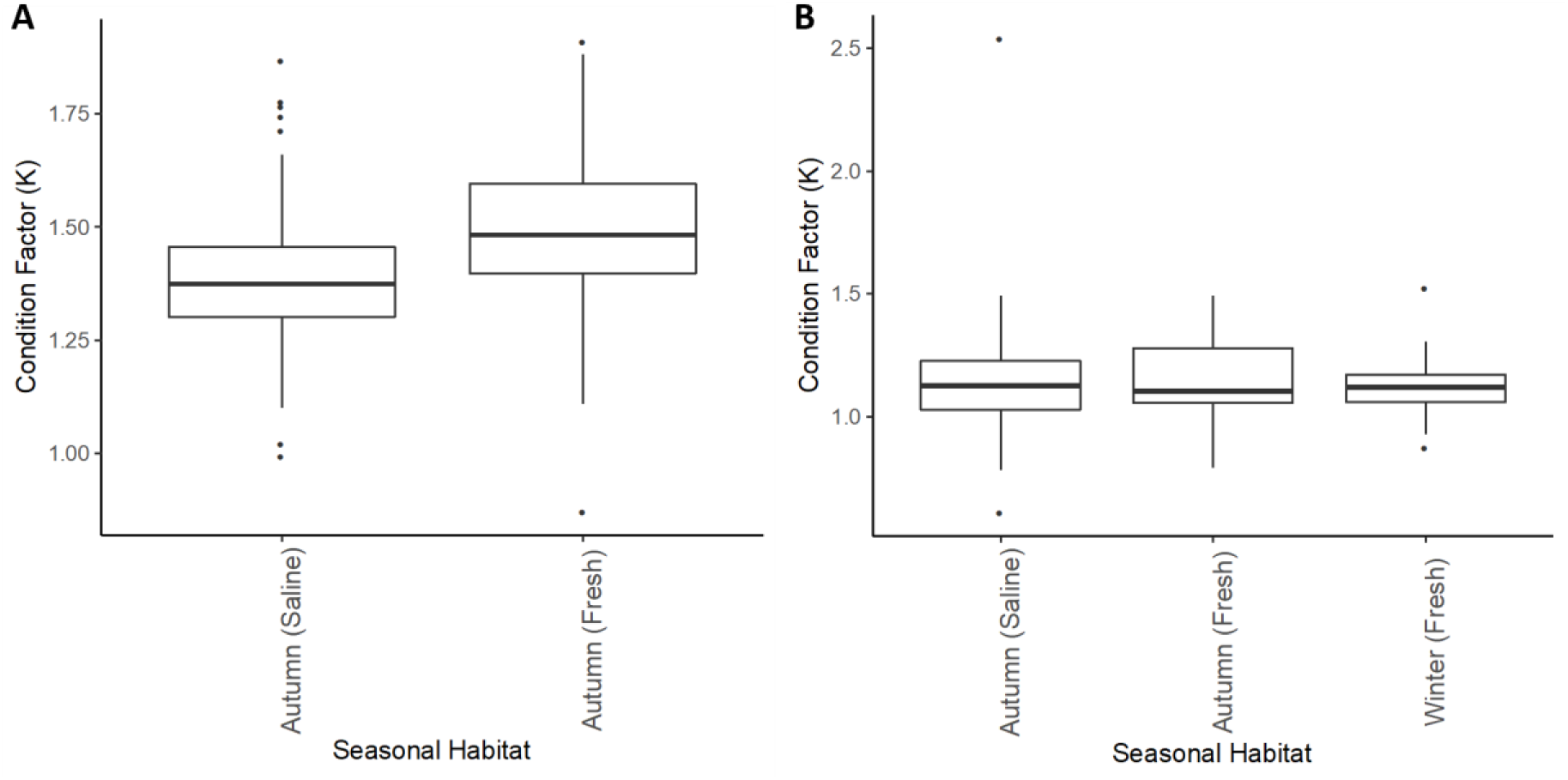
Condition factor (K) across seasonal habitat for **A)** lake whitefish where K in the autumn-fresh seasonal habitat was significantly higher (*p* < 0.001, one-way ANOVA) that that of the autumn-saline habitat and **B)** cisco where there was no significant difference between habitats.

## Supplemental Tables

**Table S1.** List of intestine-associated contaminant ASVs identified by Decontam using the prevalence method and 0.1 threshold. ASVs are sorted by score statistic value (p prev) with smallest scores indicating higher prevalence in negative controls.

| ID | p prev | Taxonomy (showing lowest taxonomic rank) |
| --- | --- | --- |
| dabbc27c1eead6a389cdf26730166d3f | 3.97E-08 | g__Allorhizobium-Neorhizobium-Pararhizobium-Rhizobium |
| 5381ac9318e2e96beb3d4919b485a2e9 | 1.31E-06 | g__Escherichia-Shigella |
| c736ae1707758eba2023209461b4bdf8 | 5.77E-06 | o__Enterobacterales |
| fade8172bb3db2dc3a922ae58d2338fb | 2.29E-05 | g__Escherichia-Shigella |
| 9b6a7459b2c92cee0abfc36d503488b7 | 0.000132 | g__Brevundimonas |
| e2c81e72268af19815d788966a4b700f | 0.000138 | <u>g</u> *Mycoplasma* |
| 4be27b69ced696653ec16ccd9b022aed | 0.000902 | <u>g</u> *Escherichia-Shigella* |
| 33bf14bcef62db17b4ae689e948f3b09 | 0.001756 | <u>g</u> *Psychrobacter* |
| ac164649e33237f799c1ffee4f344d5d | 0.002096 | <u>g</u> *Escherichia-Shigella* |
| 6914dd63f587c01fcc871d90ac6f57d7 | 0.002967 | <u>g</u> *Pseudomonas* |
| 0b24da8e753303d556f9e2dfb50a3828 | 0.012694 | <u>s</u> *Sphingomonas_echinoides* |
| 9c62ed64e515b3e0bb487790326040ff | 0.018694 | <u>s</u> *Shewanella_frigidimarina* |
| 50e0c9e86561addcca36ce6ecc4c6106 | 0.019084 | <u>g</u> *Mycoplasma* |
| 998ffc437cf8df3c16bb797dc727477f | 0.019084 | <u>g</u> *Mycoplasma* |
| a7e99d2c11263f813038356f914d126a | 0.026052 | <u>g</u> *Mycoplasma* |
| c30a5a653f7d507c84f4b23fa2a34b54 | 0.026515 | <u>s</u> *Brochothrix_thermosphacta* |
| f39be8a259561fc18be25a434936e63f | 0.038161 | <u>g</u> *Rubripirellula* |

**Table S2.**
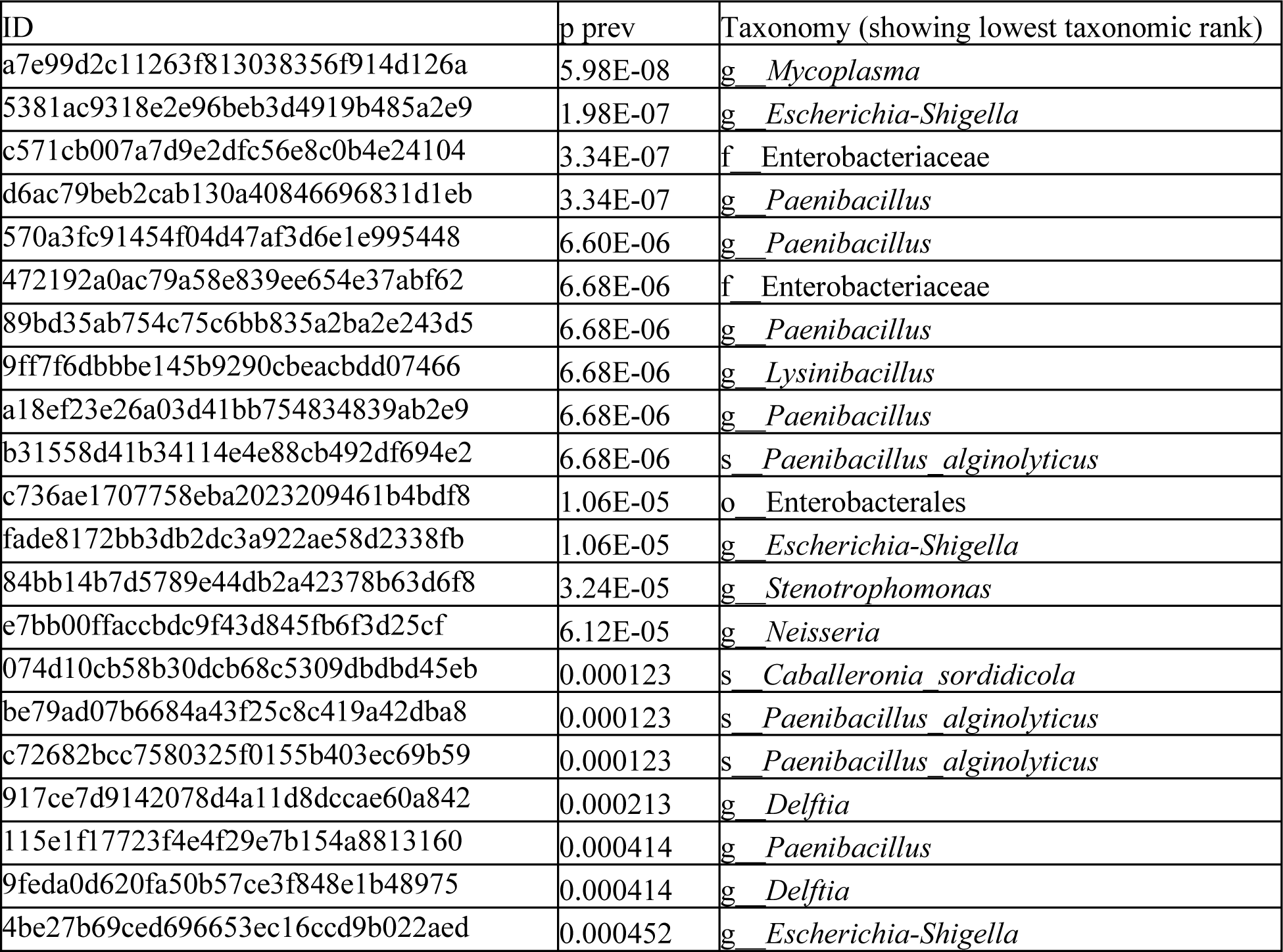
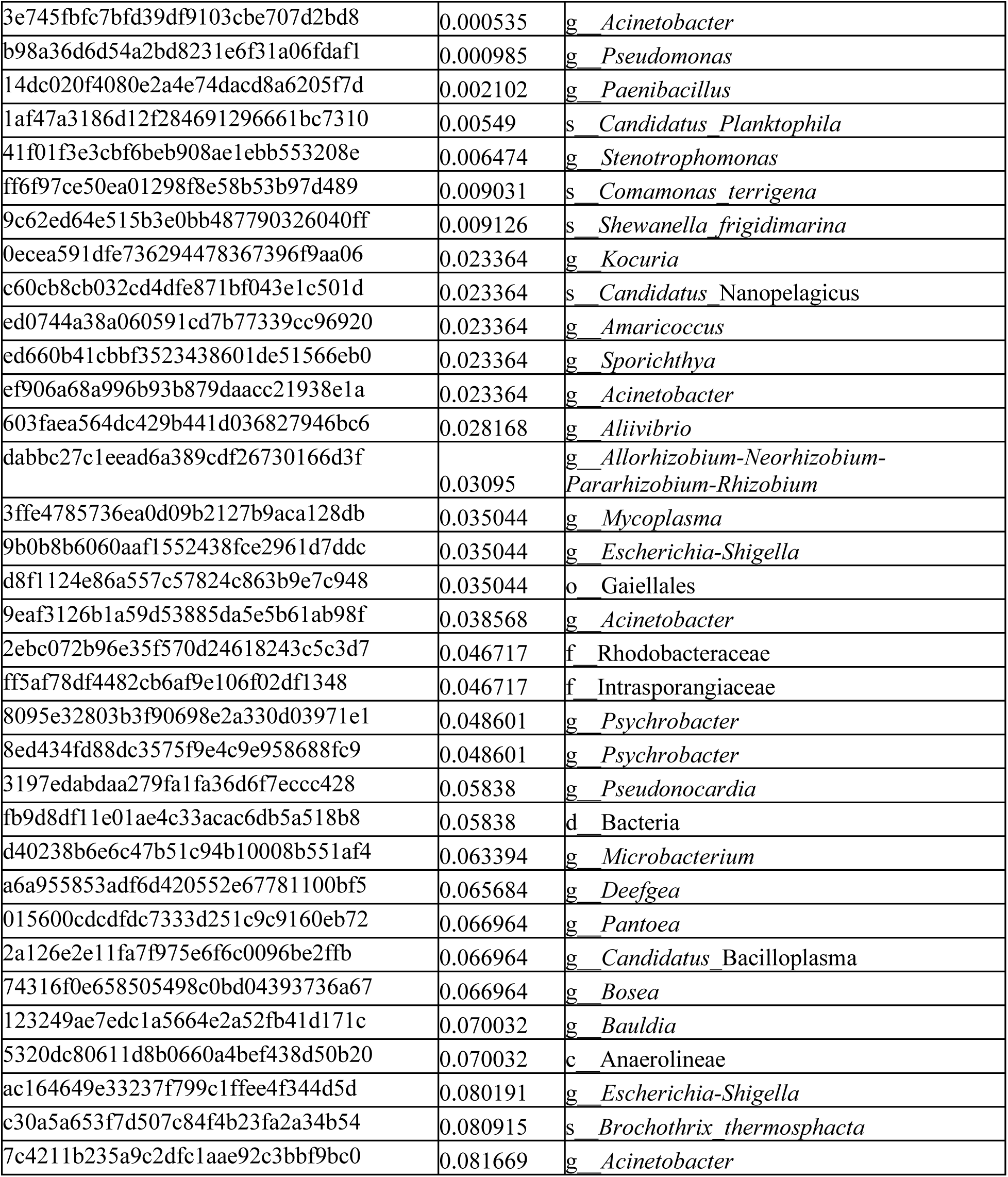
List of skin-associated contaminant ASVs identified by Decontam using the prevalence method and 0.1 threshold. ASVs are sorted by score statistic value (*p* prev) with smallest scores indicating higher prevalence in negative controls.

